# Trachomatous Trichiasis (TT) management in Tanzania: a mixed method study investigating barriers and facilitators to obtaining treatment

**DOI:** 10.1101/2020.03.13.990424

**Authors:** Rebecca M. Flueckiger, George Kabona, Upendo Mwingira, Alistidia Simon, Jeremiah Ngondi

## Abstract

**Background:** Prolonged ocular Chlamydial infection, known as trachoma, can lead to trachomatous trichiasis (TT). TT is the stage of trachoma where the eyelid turns inwards, resulting in lashes rubbing against the cornea. TT can damage the cornea, leading to vision impairment or blindness. Treatment for TT includes epilation or surgery. Trachoma is targeted for elimination as a public health problem. One criterion of trachoma elimination is less than 0.2% prevalence of TT unknown to the health system in adults >= 15 years. There are several districts in Tanzania that have not attained this target.

**Methodology:** We selected six districts across three regions in Tanzania. Our mixed-methods approach included a retrospective review and analysis of program data and implementation of key informant interviews (KII) and focus group discussions (FGD).

The desk review collated data on district-level indicators and generated estimates around number and proportion of cases not identified by case finders and cases lost along the continuum of care. KIIs and FGDs guides were structured to enlist responses around case finding techniques, linkage to services and TT surgery process.

**Conclusion:** We found a substantial proportion (13%) of TT positive people were not being identified by case finders, and of those identified, majority (72%) were lost along the continuum of care. These factors likely contribute to high TT prevalence in districts where surgical interventions are ongoing. Engaging community leaders to share TT information and enlisting people who have received surgery to witness in communities may encourage consent of examination by case finders and increase surgical uptake. After witnessing positive effects of surgery, many interviewees who had previously declined surgery changed their mind. Increasing frequency of surgical camps would improve access to these populations. Additionally, giving more notice about surgical camps and extending duration is important to enable remote populations to obtain services.

**Author Summary:** Treatment for trachomatous trichiasis (TT) includes epilation or surgery. There are several districts in Tanzania that have struggled to link people with TT to services. It is important for the program to understand why this is the case to inform program adaptations for improved linkage to services. We implemented a mixed methods approach to address this knowledge gap. We found a large portion of TT positive people are not being identified by case finders and of those identified, many are lost along the continuum of care. These factors are likely contributing to the unexpectedly high TT prevalence in districts where surgical interventions are ongoing. Barriers to identifying cases included remoteness, case finder credibility, knowledge of TT, and case finder motivation. Once cases are identified, the largest gap along the continuum of care is the link between being identified and screened. We found barriers to attending screenings and subsequently obtaining treatment to be fear of surgery, distance from surgical camps, agricultural season, time to plan, awareness and frequency of camps, and lack of assistance after surgery.

## Introduction

Prolonged conjunctival infection with *Chlamydia trachomatis* leads to an inflammatory response, trachomatous inflammation–follicular (TF) and trachomatous inflammation-intense. Overtime, cycles of repeated infection can progress to scarring of the conjunctiva, causing entropion inward turning of the eyelid and resulting in lashes rubbing against the cornea. This painful stage of the disease is called trachomatous trichiasis (TT). TT can damage the cornea, leading to vision impairment or blindness [1, 2]. Trachoma is targeted for elimination as a public health problem[3]. One criterion of trachoma elimination is prevalence of TT unknown to the health system of less than 0.2% in adults 15 years and older [4]. “Unknown to the health system” are cases that have not previously been operated on, not previously refused treatment, or have not been referred for treatment [5]. District-level prevalence is estimated through population based prevalence surveys [6] and these prevalence estimates are used as a guide for planning interventions [7, 8]. Studies have shown that TT surgery results in improved vision and physical function [9] as well as reduced photophobia and pain [10]. The World Health Organization (WHO) recommends a bilamellar tarsal rotation procedure to correct entropion caused by TT [11]. While not recommended by WHO, epilation is commonly practiced to manage minor TT and may lower risk of corneal opacity [12, 13]. Regardless of vision loss, untreated TT has been shown to significantly reduce quality of life [14, 15].

Linking TT positive people to TT surgery is a major concern in trachoma endemic settings. In Tanzania, TT surgery is provided free of charge and surgery programs are active in six regions (Mtwara, Pwani, Dodoma, Arusha, Manyara, and Lindi). The continuum of care for TT is the following: 1) Case finders identify a positive TT case, 2) Eye care professionals screen the identified case, 3) TT surgeons confirm the screened case, and 4) Treatment is provided.

In 2016, the TT backlog in Tanzania was estimated to be 214,800 [16], and in 2017, only 2,120 people were reported to have received surgery [17]. This equates to less than 1% of people needing surgery receiving it. In 2018 surveys were conducted in Mtwara, Dodoma, and Lindi which demonstrated continued TT prevalence above the elimination threshold. It is important for the trachoma programs to understand why TT elimination targets are not being achieved to inform program adaptations for improving access to services. We implemented a mixed methods approach to better understand why trachoma impact surveys in Tanzania are demonstrating unexpectedly high prevalence of TT in communities where TT surgical intervention is ongoing and how this challenge could be addressed.

## Methods

Our conceptual framework is a mixed methods approach which includes a retrospective review and analysis of program data and implementation of key informant interviews (KII) and focus group discussions (FGD).

### Sample selection

The regions of Dodoma, Lindi, and Mtwara demonstrated different levels of success in achieving TT elimination thresholds. We selected two districts in each of the three regions. The number of surgeries performed in each of these districts (in 2018 for the districts in Dodoma and Lindi and in 2016 for the districts in Mtwara) was used as an indicator for selecting districts to include in this study. We selected districts with the highest number of surgeries, as conducting many surgeries suggests that the program was active for an entire year (or close to it) rather than only conducting a few surgical camps over the course of one or two months. The districts selected were Bahi and Chamwino in Dodoma, Liwale and Ruangwa in Lindi, and Newala and Tandahimba in Mtwara.

The review of program data collated information on 8,834 people from case search registries, screening registries, TT surgery logs, and supervision reports across the six districts (Table 1). The sample was not meant to be generalizable, but rather to provide insight into the situation within these specific geographies as a supplement to the KIIs and FGDs.

**Table 1.**
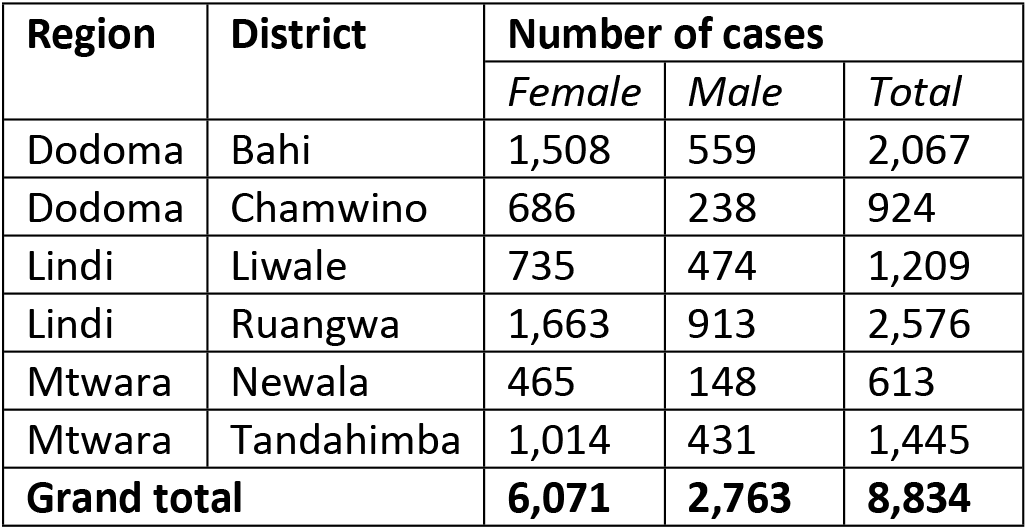
Program data sample.

| Region | District | Number of cases |  |  |
| --- | --- | --- | --- | --- |
|  |  | Female | Male | Total |
| Dodoma | Bahi | 1,508 | 559 | 2,067 |
| Dodoma | Chamwino | 686 | 238 | 924 |
| Lindi | Liwale | 735 | 474 | 1,209 |
| Lindi | Ruangwa | 1,663 | 913 | 2,576 |
| Mtwara | Newala | 465 | 148 | 613 |
| Mtwara | Tandahimba | 1,014 | 431 | 1,445 |
| **Grand total** |  | **6,071** | **2,763** | **8,834** |

The qualitative component of this study involved selecting TT positive people who had received surgery, TT positive people who had not received surgery, TT case finders (who are typically members of the community they work in), TT surgeons, and district-level health officers from the six selected districts. Sample size estimates for qualitative studies often rely on theoretical saturation which cannot be determined prior to beginning the study. Because of this, previously published estimates were used to inform the sample size [18, 19] (Table 2).The TT positive people and the case finders were purposively selected from the program data while considering distributions by ward. The ward was considered in order to ensure representation from different communities in the districts. A random selection of case finders for FGDs was made in each district. Each of the six districts had one active TT surgeon, and so this individual was interviewed in each district. The district-level health officers were selected based on their level of involvement during program implementation. In all districts, the district eyecare/NTD coordinators were interviewed. No incentives were provided for participating in these KIIs and FGD, and no recruited person declined to participate.

**Table 2.** Qualitative sample.

| Region | District | Persons with TT who received treatment |  |  | Persons with TT who did not receive treatment |  |  | Case finders (FGD) | Surgeons | District-level health officers |
| --- | --- | --- | --- | --- | --- | --- | --- | --- | --- | --- |
|  |  | Female | Male | Total | Female | Male | Total |  |  |  |
| Dodoma | Bahi | 7 | 6 | 13 | 7 | 3 | 10 | 10 (1) | 1 | 1 |
| Dodoma | Chamwino | 5 | 5 | 10 | 5 | 5 | 10 | 10 (1) | 1 | 1 |
| Lindi | Liwale | 5 | 6 | 11 | 5 | 4 | 9 | 10 (1) | 1 | 1 |
| Lindi | Ruangwa | 5 | 5 | 10 | 5 | 5 | 10 | 8 (1) | 1 | 1 |
| Mtwara | Newala | 6 | 4 | 10 | 5 | 5 | 10 | 8 (1) | 1 | 1 |
| Mtwara | Tandahimba | 6 | 4 | 10 | 5 | 5 | 10 | 10 (1) | 1 | 1 |
| **Grand total** |  | **34** | **30** | **64** | **32** | **27** | **59** | **56 (6)** | **6** | **6** |

The Interviewers and FGD facilitators were experienced in conducting qualitative studies and who are familiar with the cultural background of the study area and spoke Kiswahili fluently. Prior to the field work the interviewers and facilitators were trained by the local primary investigator on the protocol, interview guides and discussion guides.

### Analysis

#### Quantitative

All quantitative analyses were performed in R v3.4.4. From the program data, we first estimated the number of TT cases in each district (backlog) as;

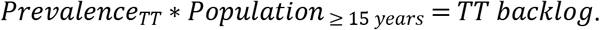

TT prevalence was provided by the NTD control program Ministry of Health, Community Development, Gender, Elderly, and Children (MoHCDGEC) (Table 3Table 3), and population estimates were calculated using methods described in detail elsewhere [16]. Briefly, district-level population was estimated from www.worldpop.org raster files [20] using the zonal statistics tool in ArcGIS 10.3[21] and then multiplied by the population percent expected to be 15 years and older derived from the UN population division (UNdata)[22] rural population pyramids.

**Table 3.** TT prevalence estimates.

| Region | District | TT prevalence | Population (15 years and older) |
| --- | --- | --- | --- |
| Dodoma | Bahi | 1.07% (2018) | 136,599 |
| Dodoma | Chamwino | 0.28%/0.47% (2018)* | 198,126 |
| Lindi | Liwale | 0.69% (2018) | 58,599 |
| Lindi | Ruangwa | 0.92% (2017) | 104,729 |
| Mtwara | Newala | 0.28% (2018) | 143,004 |
| Mtwara | Tandahimba | 0.72% (2018) | 173,056 |
| *split into North and South Chamwino evaluation units for 2018 surveys |  |  |  |

We next explored the continuum of care and determined the month each step along the continuum occurred, time lag between each step (estimating the median and interquartile range (IQR)), and proportion of cases that drop out before surgery and where along the continuum this occurred. To estimate the number of positive cases lost between being identified by case finders and screened, we assume that those who did not attend screening similar to those who did. We estimated the number of positive cases lost between identification and screening as

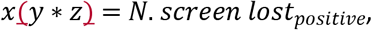

where x is percent confirmed positive, y is percent lost before screening, and z is number identified by case finders.

With these findings, we estimated the proportion of the TT backlog lost along the continuum of care by multiplying the dropout proportion at each step by the total backlog. We then calculated the number of cases not identified by case finders as

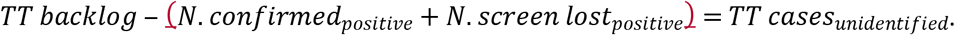

Finally, we ran regression models to determine significance of association between cases being screened and district, sex, and month of identification, as well as association between cases treated and district. The quantitative estimates are not meant to draw broad conclusions. These estimates are illustrative and give a general idea of the magnitude of gaps along the continuum of care.

#### Qualitative

All qualitative analyses were performed in Atlas.ti v8.1.28.0. A qualitative code book was developed a priori representing key constructs. These included: case finding techniques, linking identified cases to services, and surgery process. The guides also capture information on age, employment, and time of year when surgery was offered. Interviews were audio recorded, translated into English, and transcribed verbatim. The transcribed KIIs and FGDs were organized and coded according to the code book. When new themes were identified, additional codes were added. Key themes were developed and organized around the study objectives. We have provided participant quotes to illustrate the themes.

## Ethics

This study was approved by the National Institute for Medical Research of Tanzania (reference NIMR/HQ/R.8z/Vol.IX/3129). Prior to the KIIs and FGDs, a written consent was obtained from each participant. The consent form was written in Kiswahili, the local language, and Each participant was further clarified on the purpose of the study and was ensured of privacy and confidentiality. As part of the consent process, we informed participants they could end the interview at any time or refuse to answer any question. Personal identifiers were removed from all datasets before analyses were undertaken.

## Results

Here we first present the quantitative findings and then the themes identified during the qualitative component of this study.

### Quantitative findings

We estimated current backlog across the six districts to be **5,229** people, over half of which are expected to be found in Bahi and Tandahimba.

**Table 4.** District level backlog estimates.

| Region | District | TT prevalence | Population (15 years and older) | Backlog estimate |
| --- | --- | --- | --- | --- |
| Dodoma | Bahi | 1.07% | 136,599 | 1,462 |
| Dodoma | Chamwino | 0.38% | 198,126 | 753 |
| Lindi | Liwale | 0.69% | 58,599 | 404 |
| Lindi | Ruangwa | 0.92% | 104,729 | 964 |
| Mtwara | Newala | 0.28% | 143,004 | 400 |
| Mtwara | Tandahimba | 0.72% | 173,056 | 1,246 |
| **Grand total** |  |  |  | **5,229** |

We observed variation between the districts when exploring the timing of each step on the continuum of care. Bahi and Chamwino have active programs through the year, Liwale and Tandahimba for half the year, and Ruangwa and Newala for four months out of the year. In Tanzania, dry season is June through November. Here we see that 59% of cases were identified during dry season in Bahi, 40% in Chamwino, 64% in Liwale, 0% in Ruangwa, 100% in Newala and Tandahimba. Importantly, 53% of treatments occurred during dry season in Bahi, 39% in Chamwino, 79% in Liwale, 5% in Ruangwa, 100% in Newala and Tandahimba.

The time lag between being identified by a case finder and receiving treatment also varied by district. In general, the full continuum of care took between 19 and 25 days (Table 5). However, in Liwale it was much faster (median 5 days and IQR 3-8 days) and in Tandahimba it was much longer (median 97 days and IQR 92-109 days).

**Table 5.**
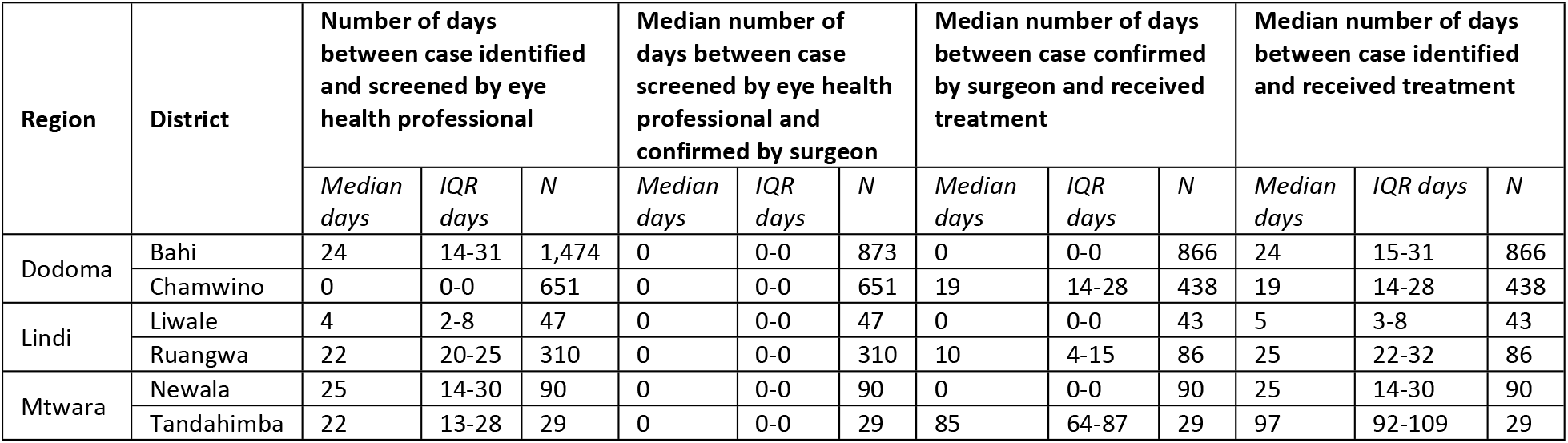
Time lag between each step along the continuum of care.

Once identified by case finders, 72% were screened by eye health professionals (ranging from 52% in Liwale to 85% in Ruangwa) (Figure 1). This means that 28% of the identified cases dropped off the continuum of care in the second step (Table 6). Of those who were recommended for treatment, 82% accepted treatment (ranging from 49% in Ruangwa to 96% in Chamwino). Of those who accepted treatment, all received treatment.

**Figure 1.**
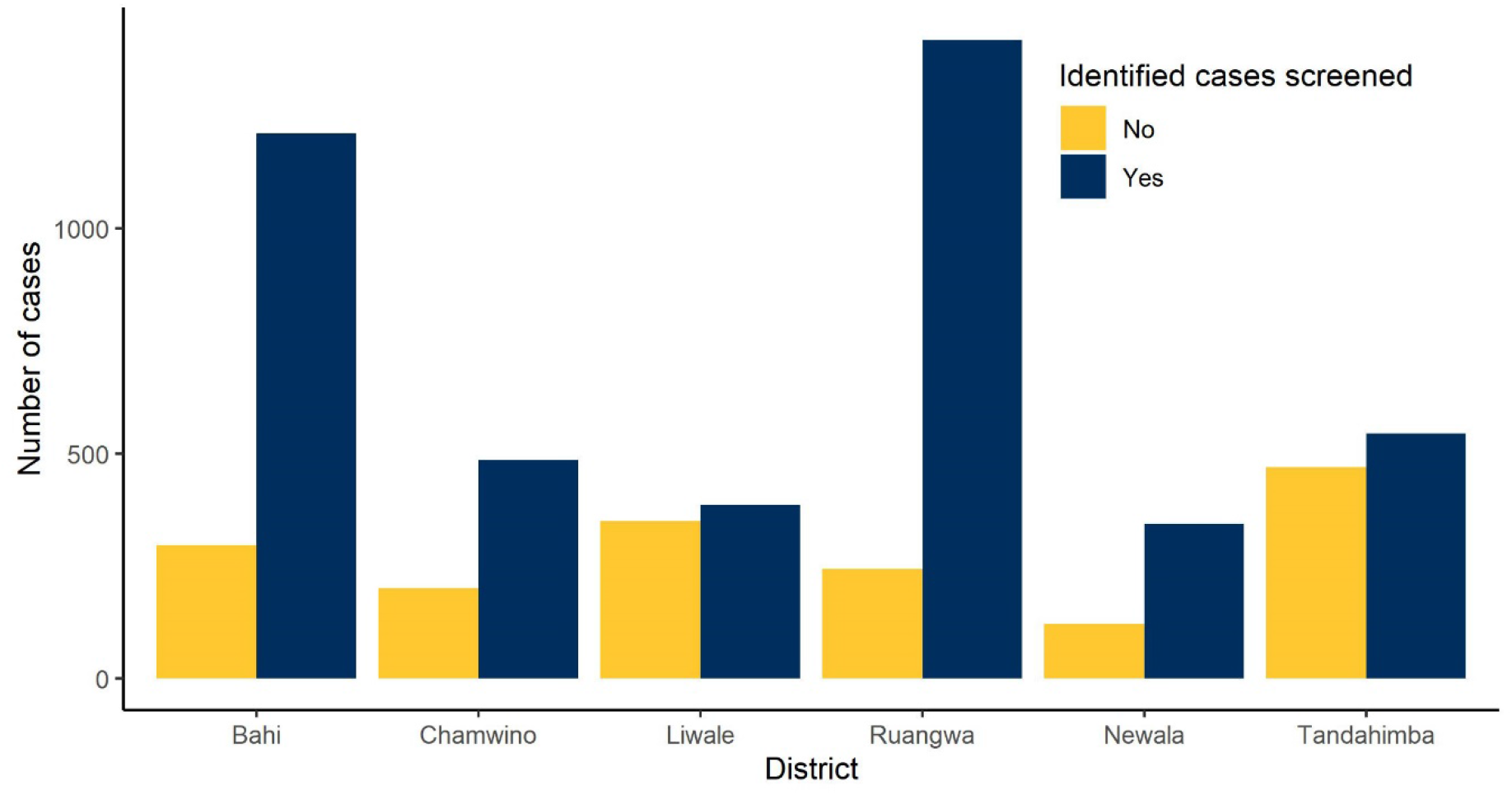
Number of identified cases screened

**Table 6.** Percent of cases along the continuum of care.

| Region | District | Percent of identified cases screened by eye health professional | Percent of screened cases confirmed as positive by eye health professional | Percent of cases confirmed by eye health professional also confirmed by surgeon | Percent of confirmed cases accepted treatment | Percent of accepted treatment cases received treatment |
| --- | --- | --- | --- | --- | --- | --- |
| Dodoma | Bahi | 79.3% | 63.2% | 100% | 95.5% | 100% |
|  | Chamwino | 71.6% | 71.7% | 100% | 96.1% | 100% |
| Lindi | Liwale | 51.8% | 23.8% | 100% | 88.6% | 100% |
|  | Ruangwa | 85.2% | 30.5% | 100% | 49.0% | 100% |
| Mtwara | Newala | 72.9% | 68.2% | 100% | 86.6% | 100% |
|  | Tandahimba | 53.8% | 75.1% | 100% | 78.8% | 100% |
| Grand total |  | 71.8% | 50.7% | 100% | 81.7% | 100% |

If we assume that those who attended screening are like those who did not, then we could estimate that across these six districts 1,327 TT positive people were lost before the second step in the continuum of care (Bahi: 270, Chamwino: 187, Liwale: 139, Ruangwa: 116, Newala: 113, and Tandahimba: 501). This means that 25% of the backlog was lost between being identified and screened (Bahi: 19%, Chamwino: 25%, Liwale: 34%, Ruangwa: 12%, Newala: 28%, and Tandahimba: 40%). We further estimate that 11% of the backlog was lost between screening and receiving treatment (Bahi: 3%, Chamwino: 3%, Liwale: 4%, Ruangwa: 35%, Newala: 10%, and Tandahimba: 10%).

We estimate that 689 cases were not identified by case finder, which equates to 13% of the backlog (Bahi: 11%, Chamwino: 13%, Liwale: 29%, Ruangwa: 18%, Newala: 0%, and Tandahimba: 13%). We further explored the dataset in an effort to identify factors that could be contributing to this disconnect. In our regression model, we found no significant association in attending screening and the month the case was identified.

The regression model where attending screening is dependent on district and sex, we found that cases identified in Bahi, Ruangwa, and Newala were significantly more likely to attend screenings and that sex was not statistically significant. The regression model where treatment acceptance was dependent on district showed positive people in Bahi were the most likely to accept treatment, followed by Chamwino, Liwale, Newala and Tandahimba. The model showed no significance for Ruangwa.

### Qualitative

Interviews were conducted among 119 people with TT across the six districts. 52% of interviewees were female (ranging from 38% to 61% across the districts). The mean (standard deviation) age of interviewed TT positive people who received surgery was 58 years (20 to 85 years old) and those who did not receive surgery was 59 years old (30-90 years old). Of those interviewed, 66% were offered surgery in the summer, 16% in the winter, 14% in the spring and 4% unknown. Of those interviewed, 93% identified themselves as “peasants”, 5% as “not employed”, and 2% “unknown”.

Below we present themes identified through the KIIs and FGDs, including feedback of case finding techniques, linkage of identified TT cases to surgeries, and the surgery process. The themes were consistent across districts and so are presented together.

#### Case finding techniques

Participants consistently identified similar processes for case finding techniques, which involved house-to-house visits where the individuals in the household were examined using a torch light and identified as having TT or not having TT. If an individual was identified as TT positive, they were provided with information on obtaining surgery. Specific themes expressed in the interviews were: case finding techniques are generally effective; knowledge about TT is a major barrier to case finding; geography is a major barrier for case finding; perceived credibility of case finders is a major barrier for case finding; and motivation of case finders is a challenge.

##### Error! Reference source not found

Nearly all who were interviewed expressed the feeling that the current case finding techniques are generally effective. People interviewed who were positive for TT specifically mentioned that they appreciated case finders coming to their home (house-to-house technique) and that this method is the best way to reach the most people.

> *“Following patients at their home is a very good technique because it will not miss patient like me who cannot walk.” – TT positive person in Tandahimba*

Case finders agreed that the house-to-house technique is an effective approach.

> *“The process is excellent, because we visit every house; this process assures that no one is left behind. If we were to invite them somewhere at central point for examination, due to various reasons some will not come.” – Case finder in Bahi knowledge about TT is a major barrier to case finding*

Most TT positive people interviewed were not aware of TT prior to being diagnosed, Though a few did mention hearing about TT on the radio or from other community member. The district-level health officers thought that this lack of knowledge may contribute to misconceptions about the program and fear of case finders.

> *“One of the barriers is poor understanding and awareness of the importance of TT surgery. When TT case finders visit the household, some people run away to avoid examination and identification.” – District-level health officer in Chamwino*

##### geography is a major barrier for case finding

The majority of those interviewed specifically mentioned that traveling far distances to remote villages is difficult for case finders and that these populations may be missed during case finding.

> *“The first challenge is the distance between households. Some are very far for the case finder to visit. The second challenge is lack of transport; the case finders have to walk from one house to the other.” – District-level health officer in Ruangwa*
>
> *“A challenge is geographical, in some communities people live far apart, we are forced to walk long distances to reach them and often we can’t make it. This is because we don’t have means of transport.” – Case finder in Liwale*

##### perceived credibility of case finders is a major barrier for case finding

Some TT positive people and surgeons interviewed mentioned that community members are uncomfortable with the case finders providing advice on eye health because they know the case finders and are not aware that they have been trained. Other TT positive persons interviewed mentioned that they felt more comfortable with the case finders because they are from their community and know and trust them. Case finders also spoke about this conflicting situation.

> *“Poor understanding of the case finder’s role by the community makes it difficult for some people to agree to an eye examination. They know and believe that a case finder is not an eye professional and even some who accept examination disagree with the examination findings.” – TT surgeon in Bahi*
>
> *“The main challenge is for the community to accept the TT case finders eye examination. They believe case finders are not eyecare professionals.” – TT surgeon in Chamwino*

##### and motivation of case finders is a challenge

District-level health officers and case finders noted that motivation of the case finders can be a challenge. In some districts the case finders are only paid for the positive cases they identify. Case finders also have other responsibilities and may find it difficult to prioritize case finding if they are not compensated in a fair way.

> *“Case finders also have their income generating activities, so if these two responsibilities are to be done at the same time the likelihood of the case finder to priorities his personal duties is high as he has to work in order to earn money.” – District-level health officer in Ruangwa*
>
> *“We case finders work as volunteers, but the task is too tough. As per now there is a very big variation when it comes to the issue of incentives to case finders. This kind of situation demoralizes us.” – Case finder in Newala*

#### Linking identified TT cases to services

Participants consistently identified similar processes when describing the steps, they took to receive surgery. It was often described that screening by eye health professionals, confirmation by surgeon and treatment happens on the same day.

> *“First, the case finder visited me at home. He examined me and identify that I have TT. He gave me the date when the TT surgery team would come to serve us. On the day of surgery, I was brought here by a case finder. They examine me again and confirmed that I have TT. I went on to receive the TT operation.” – TT positive person in Bahi*

Specific themes expressed in the interviews were all related to barriers linking people to services. The barriers identified were: fear and misconceptions; geographical distance; rainy season; insufficient time; lack of information on assistance accompanying TT surgery; and sex of TT patient.

##### fear and misconceptions

Fear was the most common barrier to obtaining surgery mentioned by all groups. It is commonly thought that TT surgery causes blindness.

> *“In some places you will find the rumors that it [TT surgery] is a blinding procedure, or very traumatic, and in other communities that the surgery is meant to destroy eyes.” – TT surgeon in Bahi*

##### geographical distance

Distance from the surgical site was also noted as a barrier among a variety of persons interviewed.

> *“There are some communities that live very far or in the areas where they can’t access health services during rainy season. Particularly those who live on the other side of the river; they cannot cross to this side.” – Case finder in Ruangwa*
>
> *“It is a very long distance from where I live to the service site. If you do not have means of transport you must walk; no one can afford to do that.” – TT positive person in Liwale*
>
> *“The difference is on the patient’s side, some need transport to reach the camp and yet they cannot afford it, so they decide to walk. These will always reach the camp very late.” – Case finder in Tandahimba*

##### rainy season

Offering surgery during rainy season may disenfranchise many people to obtain surgery. This barrier was particularly prevalent in Ruangwa.

> *“I did not receive surgery because it was a rainy season and I was busy with agricultural activities.” – TT positive person in Ruangwa*
>
> *“During rainy season people are very busy with agricultural activities. People will not accept surgery because they believe after receiving surgery, they will have to spend a number of days without working and therefore may not be able to attend their agricultural duties.” – District-level health officer in Ruangwa*
>
> *“When I was supposed to go for surgery it was a high time for agricultural activities, so I did not go because of that.” – TT positive person in Bahi*
>
> *“Many people do not receive information on availability of service, and during rainy season most people move to their farms so case finders can’t find them at their homes.” – TT positive person in Newala*
>
> *“I request the TT surgery to be offered from November onwards.” – TT positive person in Tandahimba*

##### geographical distance; rainy season; insufficient time

A common barrier mentioned was short duration of surgical camps and the lack of timely announcements about surgical camps.

> *“I did not receive surgery because when I was ready for surgery the doctors had already left, so I had nowhere to get it.” – TT positive person in Chamwino*
>
> *“The duration of service provision should be increased from one day to two or three days, without doing that we will continue making mistake by missing people.” – TT positive person in Newala*
>
> *“I waited for surgery all day without receiving it. On second day, I again waited all day without receiving surgery. When they came later for another camp, I was not in the village.” – TT positive person in Tandahimba*
>
> *“We should increase the frequency of conducting TT surgery camps. The interval needs to be small for the good of the program as well as patients whom may be lost if it takes too long to wait for the service.” – Case finder in Tandahimba*

##### ack of information on assistance accompanying TT surgery

TT positive persons interviewed were not aware of any post-operative assistance until they arrived for surgery. The need for assistance is a barrier to seeking care, and participants suggested that providing information on assistance may have encouraged some of those interviewed who did not receive surgery to receive it.

> *“They did not mention any assistance to be provided to me.” – TT positive person in Bahi (refused surgery)*
>
> *“I was not informed about any form of assistance.” – TT positive person in Chamwino (refused surgery)*
>
> *“No assistance was offered. I needed food and transport fare to go to the hospital.” – TT positive person in Tandahimba (refused surgery)*

##### ex of TT patient

In these communities it is common for men to be decision-makers, so women may have an additional barrier of convincing their husband to allow them to seek surgery.

> *“Men in our community are decision makers so when they are afraid of surgery, they will find a way to make a wife not receive surgery.” – TT positive person in Bahi (male)*
>
> *“When the sick person is a wife; the husband may refuse on her behalf.” – Case finder in Liwale*
>
> *“For women who are married, a husband decides whether his wife can go for surgery or not. Particularly during rainy season most men will not allow their wife to go for surgery, they will need them to work together in the farm.” – TT positive person in Ruangwa (female)*

#### Surgery process

Participants consistently identified similar processes for receiving surgery which involved traveling to the surgical site, being re-examined, provided surgery and then being escorted home.

> *“They queued us for examination one after the other. They told me one eye has TT, and some of my colleague had both eyes having TT. After that we queued again for surgery where they were calling one by one to the operating room. After surgery I was transported back home.” – TT positive person in Chamwino*

Specific themes expressed in the interviews were: persons with TT who received surgery were generally pleased with the outcome; conflicting opinions on assistance after surgery; and inconsistent assistance was provided after surgery.

##### ersons with TT who received surgery were generally pleased with the outcome

The majority of those interviewed specifically mentioned positive outcomes from their surgery.

> *“I had severe photophobia after I was identified I got treated and now am fine.” – TT positive person in Liwale*
>
> *“It has helped me to gain my good healthy eyes; no more pain, tearing, difficult in reading letters or even newspapers. Now I have no problem.” – TT positive person in Bahi*
>
> *“Am now fine I can do all my duties comfortably.” – TT positive person in Tandahimba*

##### conflicting opinions on assistance after surgery

There was inconsistent messaging on whether the assistance provided after surgery was sufficient. Some TT positive persons interviewed expressed appreciation for the assistance provided and felt that it was enough.

> *“It is a good assistance and is enough even for other patients.” – TT positive person in Bahi*

While others had different experiences and circumstances at home. Participants mentioned many instances where additional assistance is needed, such as patients who live alone or who are the caregivers to others.

> *“If you live alone or you have small kids to take care of and you agree to receive surgery; they will cover your eyes and you cannot do anything. There will be no one to serve my kids, so do you want my kids to die of hunger because of this surgery? – TT positive person in Ruangwa (declined surgery)*
>
> *“I will not be able to work and so will have no income for same days; and considering that I live alone this is a challenge. Who will take care of me at home after surgery? Because I know after surgery we are not allowed to cook in a smoky kitchen and sometimes both of your eyes are covered. You can’t do anything without support.– TT positive person in Ruangwa (declined surgery)*
>
> *“TT persons taking care of others at home will not accept surgery. I know some TT persons who have not received surgery because of this but they show obvious need of surgery.” – Case finder in Bahi*
>
> *“Additional assistance is needed to support those who are independent and live alone. After surgery they cannot do anything for few days therefore, they need financial assistance to run their daily life at home.” – Case finder in Bahi*
>
> *“There is an economic barrier where people who receive surgery struggle to survive for those few days postoperatively when they are not allowed to work. This makes some people decline surgery trying to avoid this challenge.” – TT positive person in Newala*

##### inconsistent assistance was provided after surgery

Many TT positive persons interviewed mentioned receiving; transportation, medicine and food after surgery. This was most common in Dodoma region. However, many others did not receive these assistances (particularly in Lindi region and Newala district).

> *“I received transportation back home after surgery and food at campsite. Medicines, postoperative care on day one for bandage removal and more medicine after bandage removal – TT positive person in Chamwino*
>
> *“No assistance was offered; I just went back home.” – TT positive person in Ruangwa*
>
> *“I only got medicine after surgery, but to go home I had to find somebody on my own to escort me back home. After I received surgery there was nothing, not even drinking water.” – TT positive person in Liwale*
>
> *“No assistance was offered to me. Even the medicine I bought myself. They said they were supposed to give medicine for free, but they forgot to come with it.” – TT positive person in Newala*

## Discussion

Linking TT positive people to TT surgery is essential for reaching the trachoma elimination targets [4]. Our findings suggest that a large portion of TT positive people are not being identified by case finders, and those who are identified are not sufficiently attending screenings and progressing along the continuum of care to receive surgery. Of those who do receive surgery, our findings show self-reported improvement in vision, relief of pain and general positive perceptions of their outcome. This is consistent with a previous study in Tanzania, which found high TT surgical success rates [23].

Our qualitative findings suggest that the current house-to-house case finding technique is an effective approach in identifying TT positive people. However, the program data illustrates that an estimated 13% of cases were not identified by case finders. Many factors are likely contributing to case finders not identifying these people. Most participants who had TT were unaware of TT prior to being diagnosed. This lack of education about TT, availability of surgery, and what the surgery entails may contribute to refusal of examination and therefore not entering the continuum of care at all. Additionally, people who live in remote locations may not be reached by case finders. Case finders are generally unpaid and do not have access to transportation, which means that they may lack the motivation and ability to reach these far away populations. An important challenge case finders face is credibility within their communities. Some people are uncomfortable with case finders providing advice on eye health and so decline to be examined. Finally, the program data shows that in some districts case finding activities take place during rainy season. This is agricultural season in Tanzania when people with agricultural responsibilities are likely to be away from their homes and so will not have an opportunity to be identified.

Once identified, the largest gap in the continuum of care is between being identified and screened. Screening by eye health professions, confirmation by surgeons, and receipt of surgery often happen on the same day. This means that when people arrange to go in for screening, they also are expecting to receive surgery. Our qualitative findings provide many barriers for screening attendance and subsequently surgical uptake. The time lag between identification and screening may play a role in attendance. People were significantly more likely to attend screenings in Bahi, Ruangwa, and Newala districts. On average, these three districts allowed nearly three weeks for an identified person to prepare themselves to attend screening. The median number of days between identification and screening was 24 in Bahi (interquartile range (IQR): 14-31), 22 in Ruangwa (IQR: 20-25) and 25 in Newala (IQR: 14-30). This aligns with our qualitative findings where interviewees mentioned needing more time to plan travel to receive surgery and arrange for help after surgery.

After surgery, patient’s eyes are covered with a dressing, and this dressing stays in place until the next day. This impedes vision and requires that someone assist the patient at home. Patients are also advised to stay away from cooking fires for the first few days after surgery. Patients who live alone or who are the caregivers to others many require a provision of food for several days after surgery. This finding is consistent with past studies where requiring help after surgery was found to be a barrier for obtaining surgery [24, 25]. As with identifying cases, lack of knowledge and misconceptions lead to fear of surgery and in turn lack of screening attendance. Additional studies have consistently found misconceptions and fear to be barriers to surgical uptake [25-27]. Logistics certainly play a role in attendance. People who have agricultural responsibilities may not be able to attend during rainy season or people who live in remote communities may not be able to travel to the surgical site. Communities that live far from central sites often do not receive information about the surgical camps until it is too late.

Engaging with community leaders to share information about TT with their communities and enlisting people who have already received surgery to witness in their communities may encourage consent of examination by case finders and increase surgical uptake. A previous study in Tanzania found village based promotion efforts [28] and witnessing others who had received surgery [29] improved surgical uptake. Other programs could be avenues for providing information on TT. Lymphatic Filariasis Mass Drug Administration campaigns, mosquito net distributions and family planning activities were all mentioned as opportunities to sensitize and distribute information on TT. Finally, providing support to case finders could improve motivation and reach of the activities.

## Acknowledgments

We would like to acknowledge to hard work of the data collection teams and the valuable input of the community advisory board. We would also like to thank the study participants for being willing to discuss their experience with TT with us and provide honest and candid feedback.

